# Low vision impairs implicit sensorimotor adaptation in response to small errors, but not large errors

**DOI:** 10.1101/2022.01.03.474829

**Authors:** Jonathan S. Tsay, Steven Tan, Marlena Chu, Richard B. Ivry, Emily A. Cooper

**Affiliations:** Department of Psychology, University of California, Berkeley; Helen Wills Neuroscience Institute, University of California, Berkeley; Herbert Wertheim School of Optometry and Vision Science, University of California, Berkeley

## Abstract

Successful goal-directed actions require constant fine-tuning of the motor system. This fine-tuning is thought to rely on an implicit adaptation process that is driven by sensory prediction errors (i.e., where you see your hand after reaching versus where you expected it to be). Individuals with low vision experience challenges with visuomotor control, but whether they also experience challenges with implicit adaptation is unknown. To explore this question, we assessed individuals with low vision and matched controls with normal vision on a visuomotor task designed to isolate implicit adaptation. We found that low vision was associated with attenuated implicit adaptation only for small visual errors, but not large visual errors. This result not only underscores an unappreciated motor learning impairment associated with low vision, but also highlights an important constraint on how low-fidelity visual information is processed by the nervous system to enable successful implicit adaptation.

**New and Noteworthy:** Whether implicit adaptation is also impacted by visual uncertainty intrinsic to the nervous system remains unknown. To test this, we examined 20 people who live with visual uncertainty in daily life due to low vision on a visuomotor task that isolates implicit adaptation. We found that low vision attenuates adaptation in response to small errors only, paving the way for new developments in rehabilitation and assistive devices for individuals with sensory impairments.

## Introduction

Our ability to enact successful goal-directed actions derives from multiple learning processes (1–7). Among these processes, implicit motor adaptation ensures that the sensorimotor system remains well-calibrated in response to changes in the body (e.g., muscle fatigue) and in the environment (e.g., a heavy jacket). This adaptive process is driven by sensory prediction errors – the difference between the predicted feedback from a motor command and the actual sensory feedback (8,9).

A spate of previous research suggests that sensory prediction errors drive implicit motor adaptation in a Bayes-optimal fashion. According to the Bayesian integration hypothesis, the learning rate for adaptation is based on a weighted signal composed of the visual feedback and a prior expectation based on a feedforward prediction (10–12). Uncertainty in the feedback, either from temporal delay (13,14) or spatial variability (10) reduces the system’s sensitivity (i.e., learning rate) to the feedback signal and, as such, reduces the extent of implicit adaptation for all error sizes (10,11,15–18). However, the attenuating effect of visual uncertainty on implicit adaptation has only been studied under conditions of *extrinsic* uncertainty, that is, uncertainty inherent to the visual feedback in the environment. Whether implicit adaptation is also impacted by visual uncertainty *intrinsic* to the nervous system remains unknown.

Here, we set out to evaluate the impact of intrinsic visual uncertainty on implicit adaptation. To test this, we assessed individuals with low vision and matched controls on visuomotor tasks designed to isolate implicit adaptation. We reasoned that low vision – defined as an impairment in visual function that interferes with visual perception and motor control in daily life (38,40–46) – induces intrinsic visual uncertainty via impairments to the eye or visual pathways (e.g., macular degeneration, glaucoma, retinitis pigmentosa, ocular albinism). Guided by the Bayesian integration hypothesis, we predicted that individuals living with low vision would therefore exhibit attenuated implicit adaptation compared to their matched controls.

## Materials and Methods

### Ethics Statement

All participants gave written informed consent in accordance with policies approved by the Institutional Review Board (Protocol Number: 2016-02-8439). Participation in the study was in exchange for monetary compensation.

### Participants

Individuals with impaired visual function that interferes with the activities of daily life (i.e., low vision) were recruited through Meredith Morgan Eye Center and via word of mouth. Participants were screened using an online survey and excluded from participation if their self-reported visual acuity in their best-seeing eye was better than 20/30 (i.e., 0.2 logMAR). Participants also reported whether they had difficulty seeing road signs: Specifically, participants were prompted with a Likert scale from 1 (very blurry) to 7 (very clear). This functional measure correlates with visual acuity (*R* = -0.5, *p* = 0.03). We also obtained survey information to determine if participants’ low vision was related to peripheral vision loss and/or central vision loss, and whether their condition was congenital or acquired (Table S1).

Twenty participants with low vision were recruited. Each participant completed two sessions that were spaced at least 24 hours apart to minimize any savings or interference (19–21). This amounted to a total of 40 online test sessions, with each session lasting approximately 45 minutes. Twenty control participants were also recruited via Prolific, a website for online participant recruitment. All control participants also completed two sessions, which amounted to 40 online test sessions, each lasting approximately 45 minutes. Participants on Prolific have been thoroughly vetted through a screening procedure to ensure data quality. Two sessions from the control data were incomplete due to technical difficulties and thus not included in the analyses.

Control participants were recruited to match the group of participants with low vision based on age, sex, handedness, and years of education. The low vision and control groups did not differ in age (*t*_36_ = −0.6, *p* = 0.58, *µ* = −3.3, [−15.4, 8.8], *D* = 0.2), handedness (*χ*^2^(1) = 0, *p* = 0.93), gender (*χ*^2^(1) = 6.1, *p* = 0.05), or years of education (*t*_3_ = −0.1, *p* = 0.89, *µ* = −0.1, [−1.5, 1.3], *D* = 0). As expected, the low vision group reported significantly more visual impairments compared to the control group based on their self-reported difficulty with reading road signs (*t*_25_ = 16.0, *p* < 0.001, *µ* = 4.6, [4.0, 5.1], *D* = 5.1).

### Apparatus

Participants used their own computer to access a webpage hosted on Google Firebase. The task was created using the OnPoint platform (22), and the task progression was controlled by JavaScript code running locally in the participant’s web browser. The size and position of stimuli were scaled based on each participant’s screen size/resolution (height = 920 ± 180 px, width = 1618 ± 433 px), which was automatically detected. For ease of exposition, the stimulus parameters reported reflect the average screen resolution in our participant population.

### Reaching task stimuli and general procedure

During the task, the participant performed small reaching movements by moving their computer cursor with their trackpad or mouse. The participant’s mouse or trackpad sensitivity (gain) was not modified, but rather left at the setting the participant was familiar with. On each trial, participants made a center-out planar movement from the center of the workspace to a visual target. A white annulus (1% of screen height: 0.24 cm in diameter) indicated the start location at the center of the screen, and a red circle (1% of screen height: 0.24 cm in diameter) indicated the target location. The radial distance of the target from the start location was 10 cm (40% of screen height). The target could appear at one of three directions from the center. Measuring angles counterclockwise and defining rightward as 0°, these directions were: 30° (upper right quadrant), 150° (upper left quadrant) and 270° (straight down). Within each experiment, triads of trials (i.e., a cycle) consisted of one trial to each of the three targets. The order in which the three targets were presented were randomized within each cycle. Note that participants with color vision deficits could still do the task since position information also indicated the difference between the start location and target location.

At the beginning of each trial, participants moved their cursor to the start location at the center of their screen. Cursor position feedback, indicated by a white dot (0.6% of screen height: 0.1 cm in diameter), was provided when the cursor was within 2 cm of the start location (10% of screen height). After maintaining the cursor in the start location for 500 ms, the target appeared at one of three locations (see above). Participants were instructed to move rapidly to the target, attempting to “slice” through the target. If the movement was not completed within 400 ms, the message “too slow” was displayed in red 20 pt. Times New Roman font at the center of the screen for 400 ms.

Feedback during this movement phase could take one of the following forms: Veridical feedback, no-feedback, or rotated non-contingent (“clamped”) feedback. During veridical feedback trials, the movement direction of the cursor was veridical with respect to the movement direction of the hand. That is, the cursor moved with their hand as would be expected for a normal computer cursor. During no-feedback trials, the cursor was extinguished as soon as the hand left the start annulus and remained off for the entire reach. During rotated clamped feedback trials, the cursor moved at a specified angular offset *relative to the position of the target*, regardless of the movement direction of the hand – a manipulation shown to isolate implicit motor adaptation (23,24). For all feedback trials, the radial position of the cursor corresponded to that of the hand up to 8 cm (the radial distance of the target), at which point, the cursor position was frozen for 50 ms, before disappearing. After completing a trial, participants moved the cursor back to the starting location. The visual cursor remained invisible until the participant moved within 2 cm of the start location, at which point the cursor appeared without any rotation.

### The impact of low vision on implicit motor adaptation for small and large errors

Participants with low vision and control participants (N = 20 per group) were tested in two sessions, with clamped feedback used to induce implicit adaptation. Numerous studies have observed that the degree of implicit adaptation saturates for visual errors greater than 5° (25,26); thus, we examined implicit adaptation in response to 3° (an error prior to the saturation zone) and 30° (an error within the saturation zone) (Figure 1a). The session order and direction (clockwise or counterclockwise) of the clamped rotation were counterbalanced across individuals. Each session consisted of 75 cycles (225 trials total), distributed across three blocks: Baseline veridical feedback block (15 cycles), rotated clamped feedback (50 cycles), and a no-feedback aftereffect block (10 cycles).

**Figure 1:**
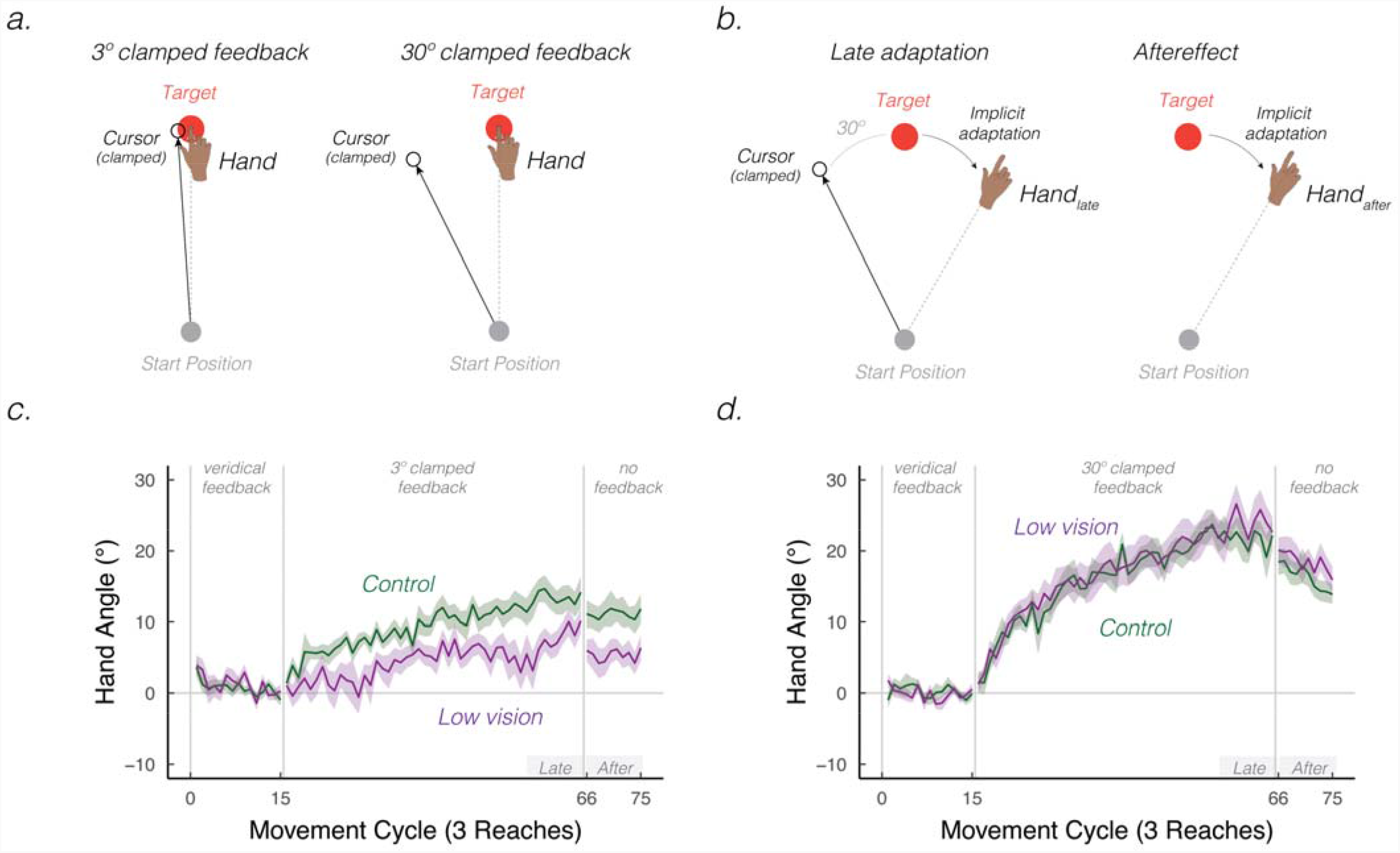
Intrinsic visual feedback uncertainty attenuates implicit adaptation in response to small, but not large errors. **(a)** The sensory prediction error – the difference between the predicted visual feedback location (i.e., the target) and perceived visual feedback location – can either be small (3°) or large (30°). **(b)** Schematic of the clamped feedback task. The cursor feedback (white circle) follows a constant trajectory rotated relative to the target (red circle), independent of the position of the participant’s hand. The rotation size remains invariant throughout the rotation block. The participant was instructed to move directly to the target and ignore the visual feedback. A robust aftereffect is observed when the visual cursor is removed during the no feedback block, implying that the clamp-induced adaptation is implicit. **(c-d)** Mean time courses of hand angle for 3° **(c)** and 30° **(d)** visual clamped feedback, for both the low vision (dark magenta) and matched control (green) groups. Hand angle is presented relative to the baseline hand angle (i.e., last five cycles of the veridical feedback block). Shaded region denotes SEM. Grey horizontal bars indicate late and aftereffect phases of the experiment.

Prior to the baseline block, the instruction “Move directly to the target as fast and accurately as you can” appeared on the screen. Prior to the clamped feedback block, the instructions were modified to read: “The white cursor will no longer be under your control. Please ignore the white cursor and continue to aim directly towards the target.” To clarify the invariant nature of the clamped feedback, three demonstration trials were provided. On all three trials, the target appeared directly above the start location on the screen (90º position) and the participant was told to reach to the left (demo 1), to the right (demo 2), and downward (demo 3). On all three of these demonstration trials, the cursor moved in a straight line, 90º offset from the target. In this way, the participant could see that the spatial trajectory of the cursor was unrelated to their own reach direction. Prior to the no-feedback aftereffect block, the participants were reminded to “Move directly to the target as fast and accurately as you can.”

### Attention and instruction checks

It is difficult in online studies to verify that participants fully attend to the task. To address this issue, we sporadically instructed participants to make specific keypresses: “Press the letter ‘b’ to proceed.” If participants failed the make an accurate keypress, the experiment was terminated. These attention checks were randomly introduced within the first 50 trials of the experiment. We also wanted to verify that the participants understood the clamped rotation manipulation. To this end, we included one instruction check after the three demonstration trials: “Identify the correct statement. Press ‘a’: I will aim away from the target and ignore the white dot. Press ‘b’: I will aim directly towards the target location and ignore the white dot.” The experiment was terminated if participants failed to make the correct response (i.e., press ‘b’).

### Data analysis

The primary dependent variable of reach performance was hand angle, defined as the angle of the hand relative to the target when movement amplitude reached an 8 cm radial distance from the start position. Specifically, we measured the angle between a line connecting the start position to the target and a line connecting the start position to the hand. Given that there is little generalization of learning between target locations spaced more than 120° apart (20,23), the data are graphed by cycles. For visualization purposes, the hand angles were flipped for blocks in which the clamp was counterclockwise with respect to the target.

Outlier responses were defined as trials in which the hand angle deviated by more than three standard deviations from a moving 5-trial window or if the hand angle was greater than 90° from the target (median percent of trials removed per participant ± IQR: control = 0.1 ± 1.0%, low vision = 0.1 ± 1.0%).

The hand angle data were baseline corrected on an individual basis to account for idiosyncratic angular biases in reaching to the three target locations (27,28). These biases were estimated based on heading angles during the last five no-feedback baseline cycles (trials 31 – 45), with these bias measures then subtracted from the data for each cycle. We defined two summary measures of learning: late adaptation and aftereffect (Figure 1b). Late adaptation was defined as the mean hand angle over the last 10 movement cycles of the rotation block (trials 166 - 195). The aftereffect was operationalized as the mean angle over all movement cycles of the no-feedback aftereffect block (trials 196 - 225).

These data were submitted to a linear mixed effect model, with hand angle measures as the dependent variable. We included Experiment Phase (late adaptation, aftereffect), Group (low vision or control), and Error Size (3°, 30°) as fixed effects and Participant ID as a random effect. A priori, we hypothesized that the low vision group would differ from the controls in their response to the small errors. Reaction time was included as a covariate in the analysis since RT was greater in the low vision group compared to the control group (RT: *t*_32_ = 3.3, *p* = 0.002, *µ* = 116.8, [47.6, 185.9], *D* = 1.1; median RT ± IQR, low vision = 425.0 ± 206.5 ms; control = 317.0 ± 133.6 ms). RT was defined as the interval between target presentation to the start of movement (operationalized as time at which the hand movement exceeded 1 cm). Movement time (MT = the time between the start of the movement and when the radial distance of the movement reached 8 cm) was not included as a covariate since MT did not differ between groups (MT: *t*_34_ = 0.6, *p* = 0.58, *µ* = 17.5, [−46.9, 81.9], *D* = 0.2; median MT ± IQR, low vision = 140.0 ± 106.5 ms; control = 106.4 ± 163.9 ms). We note that the results are similar if MT is included as a covariate.

We employed F-tests with the Satterthwaite method to evaluate whether the coefficients obtained from the linear mixed effects model were statistically significant (R functions: lmer, lmerTest, anova). Pairwise post-hoc t-tests (two-tailed) were used to compare hand angle measures between the low vision and control groups (R function: emmeans). P-values were adjusted for multiple comparisons using the Tukey method. The degrees of freedom were also adjusted when the variances between groups were not equal. 95% confidence intervals for group comparisons (t-tests) obtained from the linear mixed effects model are reported in squared brackets. Standard effect size measures are also provided (*D* for between-participant comparisons; *D*_*z*_ for within-participant comparisons; 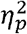 for between-subjects ANOVA) (29).

## Results

Consistent with numerous prior studies, participants in both groups showed a gradual change in hand angle in the opposite direction of the clamped feedback, trending towards an asymptotic level (Figure 1c-d) (23,35,36). We summarize these changes in hand angle relative to baseline (i.e., 0°; last five cycles of the veridical feedback baseline block) across two phases: late adaptation and aftereffect (aftereffect was measured on trials after visual feedback was removed).

Late adaptation was significant in all four conditions, indicating robust implicit adaptation generated by the clamped feedback, regardless of error size or participant vision level (3° controls: *t*_l7_ = 7.8, *p* < 0.001, *µ* = 13.4, [9.9, 17.1], *D* = 1.8; 30° controls: *t*_l7_ = 13.9, *p* < 0.001, *µ* = 21.6, [18.4, 24.9], *D* = 3.1; 3° low vision: *t*_l9_ = 5.6, *p* < 0.001, *µ* = 7.5, [4.7, 10.2], *D* = 1.3); 30° low vision: *t*_l9_ = 10.0, *p* < 0.001, *µ* = 23.8, [18.8, 28.8], *D* = 2.2). The main effect of phase was not significant (*F*_1,110_ = 1.0, *p* = 0.32, 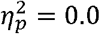), indicating that implicit adaptation exhibited minimal decay back to baseline when visual feedback was removed. Comparing the left and right panels of Figures 1c and 1d, the learning functions were higher when the error was 30° compared to when the error was 3° (*F*_1,112_ = 21.6, *p* < 0.001, 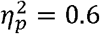), corroborating previous reports showing that implicit adaptation increases with the size of the error (26,37).

We next turned to our main question, asking how low vision impacts implicit adaptation in response sensory prediction errors. There was a significant interaction between group and error size (*F*_1,111_ = 10.5, *p* = 0.002, 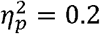): Whereas the learning function between the two groups were indistinguishable in response to a 30° error (Figure 1d; 2b), the learning function in the low vision group was attenuated compared to controls in response to a 3° error (Figure 1c; 2a). This assessment was confirmed by post-hoc t-tests, revealing that low vision was associated with attenuated implicit adaptation in response to the small error (*t*_60_ = −3.0, *p* = 0.02, *µ* = −5.8, [−9.6, −1.9], *D* = 0.9) but not the large error (*t*_57_ = 1.1, *p* = 0.67, *µ* = 2.2, [−1.6, 5.9], *D* = 0.2) (also see *Supplemental Results* for covariate analyses).

**Figure 2:**
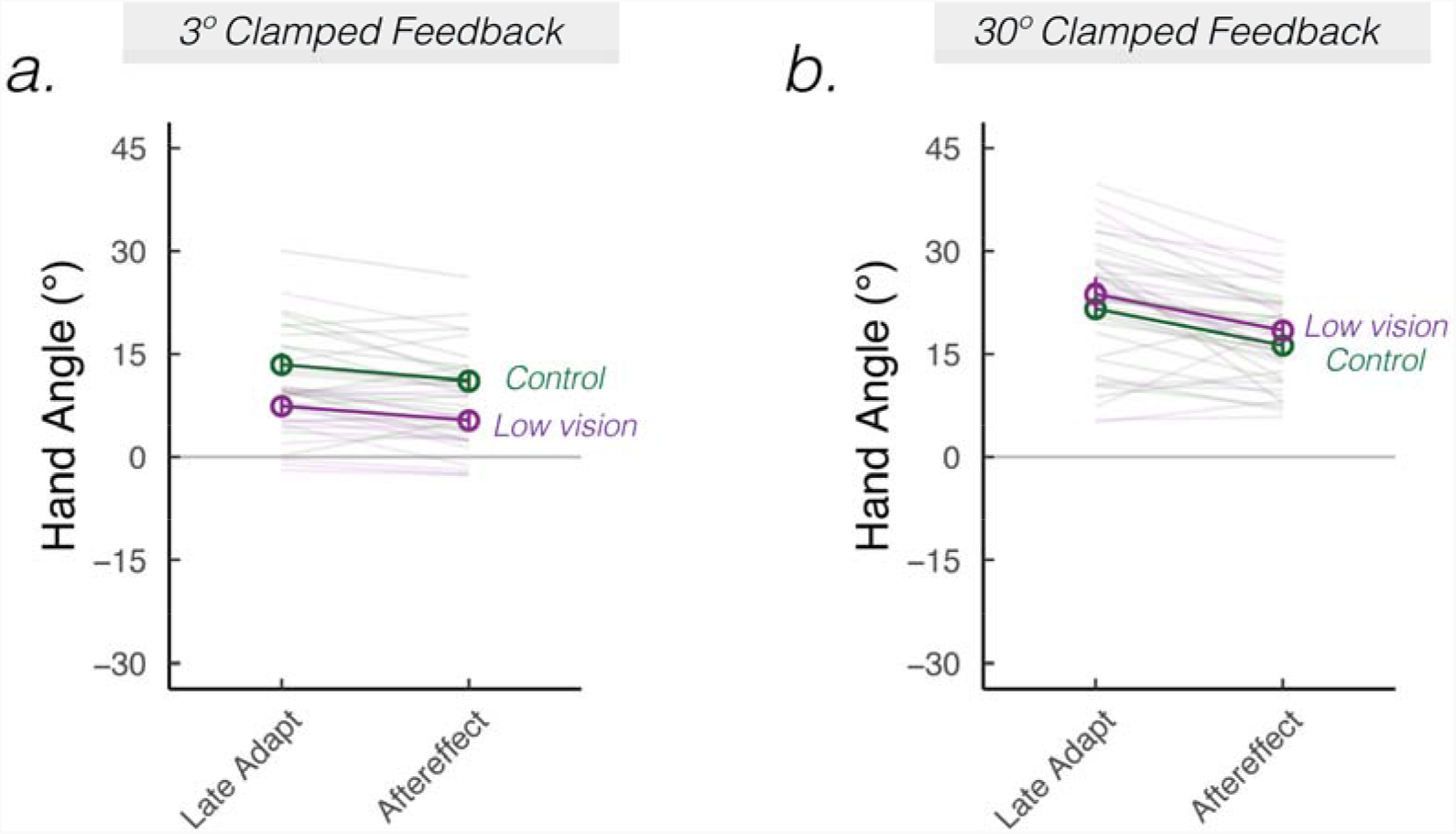
Visual uncertainty attenuates implicit adaptation in response to small, but not large errors. Mean hand angles ± SEM during early and late phases of the rotation block, and during the no-feedback aftereffect block, for 3° (**a**) and 30° **(b)** clamped rotation sessions. Thin lines denote individual participants.

Together, these results underscored an unappreciated implicit adaptation impairment associated with low vision, but only when the size of the visual error was small.

## Discussion

Low vision can cause difficulty in discriminating the position of visual objects (38,39). This impairment impacts motor performance, resulting in slower and less accurate goal-directed movements (38,40–46). Here we asked how low vision impacts *motor learning* using a visuomotor adaptation task that isolates implicit adaptation. The results revealed that low vision was associated with attenuated implicit adaptation when the sensory prediction error was small, but not when the error was large.

The error size by intrinsic visual uncertainty interaction converges with a recent in-laboratory study, in which sensory uncertainty was artificially increased using different cursor patterns (single dot = certain feedback, cloud of dots = uncertain feedback) (47). In the previous study, participants with typical vision also showed attenuated implicit adaptation with uncertain feedback when the error size was 3° but not when the error size was 30°. Together, these results point to a strong correspondence between the effect of extrinsic uncertainty in the visual stimulus (e.g., a foggy day) and intrinsic uncertainty induced by low vision (e.g., damage to or pathology of the visual system), and signal an important translation between basic and clinical science. Below, we speculate about different mechanisms that may give rise to this interaction.

A standard Bayesian integration hypothesis posits that intrinsic uncertainty induced by low vision would be associated with decreased sensitivity to errors and attenuate implicit adaptation for *all* error sizes. Therefore, the standard Bayesian integration hypothesis cannot account for our results. That being said, this error by uncertainty interaction can be accounted for by a modified Bayesian perspective, which posits that the nervous system performs causal inference (48,49): small errors, attributed to a miscalibration in the sensorimotor system, require implicit adaptation; the weight given to these small errors will fall off with increasing uncertainty. On the other hand, large errors are more likely attributed to an external source and will therefore get discounted; paradoxically, the weight given to these large errors will increase with uncertainty, since uncertainty can obscure the attribution of large errors to an external source. As such, the causal inference model predicts a crossover point, where implicit adaptation will be higher for small certain errors (compared to small uncertain errors) but then lower for large certain errors (compared to large uncertain errors).

While the null effect of visual uncertainty in our data for large visual errors supports the causal inference hypothesis, the cross-over prediction – an enhancing effect of visual uncertainty for large visual errors – was not observed (see also: (21)). A recent theory of implicit adaptation proposes an alternative possibility: The proprioceptive re-alignment hypothesis centers on the notion that implicit adaptation is driven to reduce a proprioceptive error, the mismatch between the felt and desired position of the hand, rather than a visual error (50). According to the proprioceptive re-alignment hypothesis, visual uncertainty indirectly affects implicit adaptation by influencing the magnitude of the proprioceptive shift, that is, the degree to which visual feedback recalibrates (biases) the felt position of the hand (51). When the visual error is small, visual uncertainty attenuates the size of proprioceptive shifts, and therefore, attenuates the degree of implicit adaptation. When the visual error is larger than ∼10°, proprioceptive shifts saturate and are therefore invariant to uncertainty (24,33,52). As such, visual uncertainty has no impact on implicit adaptation when the visual error is large. Our data thus offers preliminary support for the notion that implicit adaptation is driven by a process of proprioceptive re-alignment.

Previous work has shown that individuals with low vision move slower and make more errors when performing goal-directed movements (38,42,44,45,53). While these deficits are observed in people with both central and peripheral vision loss, reductions in central vision appear to be the key limiting factor (42,53). Central vision loss, which can result in lower acuity and worse contrast sensitivity, likely worsens the ability to precisely locate the intended visual target as well as respond to the sensory predictions conveying motor performance, an impairment that would be especially pronounced when the target and error are small (54,55). Interestingly, in our web-based studies we did not observe strong associations between subjective measures of visual ability and implicit adaptation (see Figure S3 c,f,i). However, future psychophysical studies recruiting specific low vision subgroups to the laboratory would be essential to home in on how different visual impairments can interact to influence implicit adaptation.

Importantly, our results suggest that individuals with low vision exhibit intact error-based learning when the error is large and clearly detected. This finding may be exploited to enhance motor outcomes during rehabilitation (56). For example, clinicians and practitioners could use non-visual feedback (e.g., auditory or tactile) to enhance the saliency and possibly reduce localization uncertainty of small visual error signals (43). Haptic wearables have the potential to provide vibration, texture, slip, temperature, force, and proprioception sensations (57), which may help to improve motor control in both reaching and grasping for individuals with low vision. Moreover, rehabilitative specialists could provide explicit instructions to highlight the presence of small errors, such that individuals may learn to rely more on explicit re-aiming strategies to compensate for these errors (58). Future work could examine which of these techniques is most effective to enhance motor learning when errors are small.

## References

1. McDougle SD, Bond KM, Taylor JA. Explicit and Implicit Processes Constitute the Fast and Slow Processes of Sensorimotor Learning. J Neurosci. 2015 Jul 1;35(26):9568–79.

2. Bond KM, Taylor JA. Flexible explicit but rigid implicit learning in a visuomotor adaptation task. J Neurophysiol. 2015 Jun 1;113(10):3836–49.

3. Taylor JA, Krakauer JW, Ivry RB. Explicit and implicit contributions to learning in a sensorimotor adaptation task. J Neurosci. 2014 Feb 19;34(8):3023–32.

4. Taylor JA, Ivry RB. Flexible cognitive strategies during motor learning. PLoS Comput Biol. 2011 Mar;7(3):e1001096.

5. Keisler A, Shadmehr R. A shared resource between declarative memory and motor memory. J Neurosci. 2010 Nov 3;30(44):14817–23.

6. Haith AM, Huberdeau DM, Krakauer JW. The influence of movement preparation time on the expression of visuomotor learning and savings. J Neurosci. 2015 Apr 1;35(13):5109–17.

7. McDougle SD, Boggess MJ, Crossley MJ, Parvin D, Ivry RB, Taylor JA. Credit assignment in movement-dependent reinforcement learning. Proc Natl Acad Sci U S A. 2016 Jun 14s;113(24):6797–802.

8. Shadmehr R, Smith MA, Krakauer J. Error correction, sensory prediction, and adaptation in motor control. Annu Rev Neurosci. 2010;33:89–108.

9. Kim HE, Avraham G, Ivry RB. The Psychology of Reaching: Action Selection, Movement Implementation, and Sensorimotor Learning. Annu Rev Psychol [Internet]. 2020 Sep 25; Available from: http://dx.doi.org/10.1146/annurev-psych-010419-051053

10. Burge J, Ernst MO, Banks MS. The statistical determinants of adaptation rate in human reaching. J Vis. 2008 Apr 23;8(4):20.1-19.

11. Wei K, Körding K. Uncertainty of feedback and state estimation determines the speed of motor adaptation. Front Comput Neurosci. 2010 May 11;4:11.

12. Körding KP, Wolpert DM. Bayesian integration in sensorimotor learning. Nature. 2004 Jan 15;427(6971):244–7.

13. Brudner SN, Kethidi N, Graeupner D, Ivry RB, Taylor JA. Delayed feedback during sensorimotor learning selectively disrupts adaptation but not strategy use. J Neurophysiol. 2016 Mar;115(3):1499–511.

14. Kitazawa S, Kohno T, Uka T. Effects of delayed visual information on the rate and amount of prism adaptation in the human. J Neurosci. 1995 Nov;15(11):7644–52.

15. Samad M, Chung AJ, Shams L. Perception of body ownership is driven by Bayesian sensory inference. PLoS One. 2015 Feb 6;10(2):e0117178.

16. van Beers RJ. How does our motor system determine its learning rate? PLoS One. 2012 Nov 12;7(11):e49373.

17. van Beers RJ, Wolpert DM, Haggard P. When feeling is more important than seeing in sensorimotor adaptation. Curr Biol. 2002 May 14;12(10):834–7.

18. Shyr MC, Joshi SS. Validation of the Bayesian sensory uncertainty model of motor adaptation with a remote experimental paradigm. In: 2021 IEEE 2nd International Conference on Human-Machine Systems (ICHMS). 2021. p. 1–6.

19. Avraham G, Morehead R, Kim HE, Ivry RB. Reexposure to a sensorimotor perturbation produces opposite effects on explicit and implicit learning processes. PLoS Biol. 2021 Mar 5;19(3):e3001147.

20. Krakauer J, Ghez C, Ghilardi MF. Adaptation to visuomotor transformations: consolidation, interference, and forgetting. J Neurosci. 2005 Jan 12;25(2):473–8.

21. Lerner G, Albert S, Caffaro PA, Villalta JI, Jacobacci F, Shadmehr R, et al. The Origins of Anterograde Interference in Visuomotor Adaptation. Cereb Cortex [Internet]. 2020 Mar 4; Available from: http://dx.doi.org/10.1093/cercor/bhaa016

22. Tsay JS, Lee A, Ivry RB, Avraham G. Moving outside the lab: The viability of conducting sensorimotor learning studies online. Neurons, Behavior, Data analysis, and Theory [Internet]. 2021 Jul 30; Available from: http://dx.doi.org/10.51628/001c.26985

23. Morehead R, Taylor JA, Parvin DE, Ivry RB. Characteristics of Implicit Sensorimotor Adaptation Revealed by Task-irrelevant Clamped Feedback. J Cogn Neurosci. 2017 Jun;29(6):1061–74.

24. Tsay JS, Parvin DE, Ivry RB. Continuous reports of sensed hand position during sensorimotor adaptation. J Neurophysiol. 2020 Oct 1;124(4):1122–30.

25. Hayashi T, Kato Y, Nozaki D. Divisively Normalized Integration of Multisensory Error Information Develops Motor Memories Specific to Vision and Proprioception. J Neurosci. 2020 Feb 12;40(7):1560–70.

26. Kim HE, Morehead R, Parvin DE, Moazzezi R, Ivry RB. Invariant errors reveal limitations in motor correction rather than constraints on error sensitivity. Commun Biol. 2018 Mar 22;1:19.

27. Morehead JR, Ivry R. Intrinsic biases systematically affect visuomotor adaptation experiments. Society for Neural Control of Movement [Internet]. 2015; Available from: http://ivrylab.berkeley.edu/uploads/4/1/1/5/41152143/morehead_ncm2015.pdf

28. Vindras P, Desmurget M, Prablanc C, Viviani P. Pointing errors reflect biases in the perception of the initial hand position. J Neurophysiol. 1998 Jun;79(6):3290–4.

29. Lakens D. Calculating and reporting effect sizes to facilitate cumulative science: a practical primer for t-tests and ANOVAs. Front Psychol. 2013 Nov 26;4:863.

30. Poh E, Al-Fawakari N, Tam R, Taylor JA, McDougle SD. Generalization of motor learning in psychological space [Internet]. bioRxiv. 2021 [cited 2021 Sep 17]. p. 2021.02.09.430542. Available from: https://www.biorxiv.org/content/10.1101/2021.02.09.430542v2

31. Hutter SA, Taylor JA. Relative sensitivity of explicit reaiming and implicit motor adaptation. J Neurophysiol. 2018 Nov 1;120(5):2640–8.

32. Liu Y, Jiang W, Bi Y, Wei K. Sensorimotor knowledge from task-irrelevant feedback contributes to motor learning. J Neurophysiol. 2021 Sep 1;126(3):723–35.

33. Tsay JS, Kim HE, Parvin DE, Stover AR, Ivry RB. Individual differences in proprioception predict the extent of implicit sensorimotor adaptation. J Neurophysiol [Internet]. 2021 Mar 3; Available from: http://dx.doi.org/10.1152/jn.00585.2020

34. Tsay JS, Haith A, Ivry RB, Kim HE. Distinct processing of sensory prediction error and task error during motor learning [Internet]. bioRxiv. bioRxiv; 2021. Available from: http://dx.doi.org/10.1101/2021.06.20.449180

35. Kim HE, Parvin DE, Ivry RB. The influence of task outcome on implicit motor learning. Elife [Internet]. 2019 Apr 29;8. Available from: http://dx.doi.org/10.7554/eLife.39882

36. Tsay JS, Haith AM, Ivry RB, Kim HE. Interactions between sensory prediction error and task error during implicit motor learning. PLoS Comput Biol. 2022 Mar;18(3):e1010005.

37. Marko MK, Haith AM, Harran MD, Shadmehr R. Sensitivity to prediction error in reach adaptation. J Neurophysiol. 2012 Sep;108(6):1752–63.

38. Timmis MA, Pardhan S. The effect of central visual impairment on manual prehension when tasked with transporting-to-place an object accurately to a new location. Invest Ophthalmol Vis Sci. 2012 May 14;53(6):2812–22.

39. Massof RW, Fletcher DC. Evaluation of the NEI visual functioning questionnaire as an interval measure of visual ability in low vision. Vision Res. 2001 Feb;41(3):397–413.

40. Cheong Y, Ling C, Shehab R. An Empirical Comparison between the Effects of Normal and Low Vision on Kinematics of a Mouse-Mediated Pointing Movement. International Journal of Human–Computer Interaction. 2021 Aug 1;1–11.

41. Lenoble Q, Corveleyn X, Tran THC, Rouland J-F, Boucart M. Can I reach it? A study in age-related macular degeneration and glaucoma patients. Vis cogn. 2019 Nov 26;27(9–10):732–9.

42. Pardhan S, Gonzalez-Alvarez C, Subramanian A. Target contrast affects reaching and grasping in the visually impaired subjects. Optom Vis Sci. 2012 Apr;89(4):426–34.

43. Endo T, Kanda H, Hirota M, Morimoto T, Nishida K, Fujikado T. False reaching movements in localization test and effect of auditory feedback in simulated ultra-low vision subjects and patients with retinitis pigmentosa. Graefes Arch Clin Exp Ophthalmol. 2016 May;254(5):947–56.

44. Kotecha A, O’Leary N, Melmoth D, Grant S, Crabb DP. The functional consequences of glaucoma for eye-hand coordination. Invest Ophthalmol Vis Sci. 2009 Jan;50(1):203–13.

45. Verghese P, Tyson TL, Ghahghaei S, Fletcher DC. Depth Perception and Grasp in Central Field Loss. Invest Ophthalmol Vis Sci. 2016 Mar;57(3):1476–87.

46. Jacko JA, Barreto AB, Marmet GJ, Chu JYM, Bautsch HS, Scott IU, et al. Low vision: the role of visual acuity in the efficiency of cursor movement. In: Proceedings of the fourth international ACM conference on Assistive technologies. New York, NY, USA: Association for Computing Machinery; 2000. p. 1–8. (Assets ‘00).

47. Tsay JS, Avraham G, Kim HE, Parvin DE, Wang Z, Ivry RB. The Effect of Visual Uncertainty on Implicit Motor Adaptation. J Neurophysiol [Internet]. 2020 Nov 25; Available from: https://journals.physiology.org/doi/abs/10.1152/jn.00493.2020

48. Wei K, Körding K. Relevance of error: what drives motor adaptation? J Neurophysiol. 2009 Feb;101(2):655–64.

49. Shams L, Beierholm UR. Causal inference in perception. Trends Cogn Sci. 2010 Sep;14(9):425–32.

50. Tsay JS, Kim HE, Haith AM, Ivry RB. Proprioceptive Re-alignment drives Implicit Sensorimotor Adaptation [Internet]. bioRxiv. 2021. Available from: http://dx.doi.org/10.1101/2021.12.21.473747

51. Cressman EK, Henriques DYP. Motor adaptation and proprioceptive recalibration. Prog Brain Res. 2011;191:91–9.

52. ‘t Hart BM, Ruttle JE, Henriques DYP. Proprioceptive recalibration generalizes relative to hand position [Internet]. 2020 [cited 2020 Dec 29]. Available from: https://deniseh.lab.yorku.ca/files/2020/05/tHart_SfN_2019.pdf?x64373

53. Pardhan S, Gonzalez-Alvarez C, Subramanian A. How does the presence and duration of central visual impairment affect reaching and grasping movements? Ophthalmic Physiol Opt. 2011 May;31(3):233–9.

54. Legge GE, Parish DH, Luebker A, Wurm LH. Psychophysics of reading. XI. Comparing color contrast and luminance contrast. J Opt Soc Am A. 1990 Oct;7(10):2002–10.

55. Tomkinson CR. Accurate assessment of visual acuity in low vision patients. Optom Vis Sci. 1974 May;51(5):321–4.

56. Tsay JS, Winstein CJ. Five Features to Look for in Early-Phase Clinical Intervention Studies. Neurorehabil Neural Repair. 2020 Nov 26;1545968320975439.

57. Patel S, Park H, Bonato P, Chan L, Rodgers M. A review of wearable sensors and systems with application in rehabilitation. J Neuroeng Rehabil. 2012 Dec;9(1):21.

58. Merabet LB, Connors EC, Halko MA, Sánchez J. Teaching the blind to find their way by playing video games. PLoS One. 2012 Sep 19;7(9):e44958.

