## Supplemental Data 1 for "Low vision impairs implicit sensorimotor adaptation in response to small errors, but not large errors"

**Supplemental Results**

Since the low vision group on average initiated movements slower than the control group (see Methods), we added RT as a covariate in our analyses. The hand angle data was not modulated by RT ($F_{1, 95}= 1.2, p=0.27,\eta_{p}^{2}=0.0$). More importantly, the error size by group interaction still holds ($F_{1, 112}= 11.4, p=0.001,\eta_{p}^{2}=0.2$), with the low vision group exhibiting attenuated implicit adaptation in response to small errors ($t_{60}= -2.4, p=0.04, \mu=-5.0, \left[ 0.9, 9.1 \right], D=0.9$), but not large errors ($t_{60}= -1.4, p=0.47, \mu=-3.1, \left[ -7.2, 1.1 \right], D= 0.2$).

We also explored whether various subgroups of the participants with low vision exhibited differences in implicit adaptation. As shown in Figure S1, there were no appreciable differences between participants with and without central vision loss (Figure S1a,b; $F_{1,18}= 0.3, p=0.62,\eta_{p}^{2}=0.0$), with and without peripheral vision loss (Figure S1c,d; $F_{1,18}= 1.0, p=0.32,\eta_{p}^{2}=0.1$), or early vs late onset of low vision (Figure S1e,f; $F_{1,17}= 0.7, p=0.51,\eta_{p}^{2}=0.1$). Furthermore, neither the participants’ self-reports of visual acuity (Figure S2a,d) nor ability to perceive road signs (Figure S2b, e) correlated with implicit adaptation. In sum, we did not identify additional features amongst individuals in the low vision group that impacted implicit adaptation.

**Supplemental Figures**

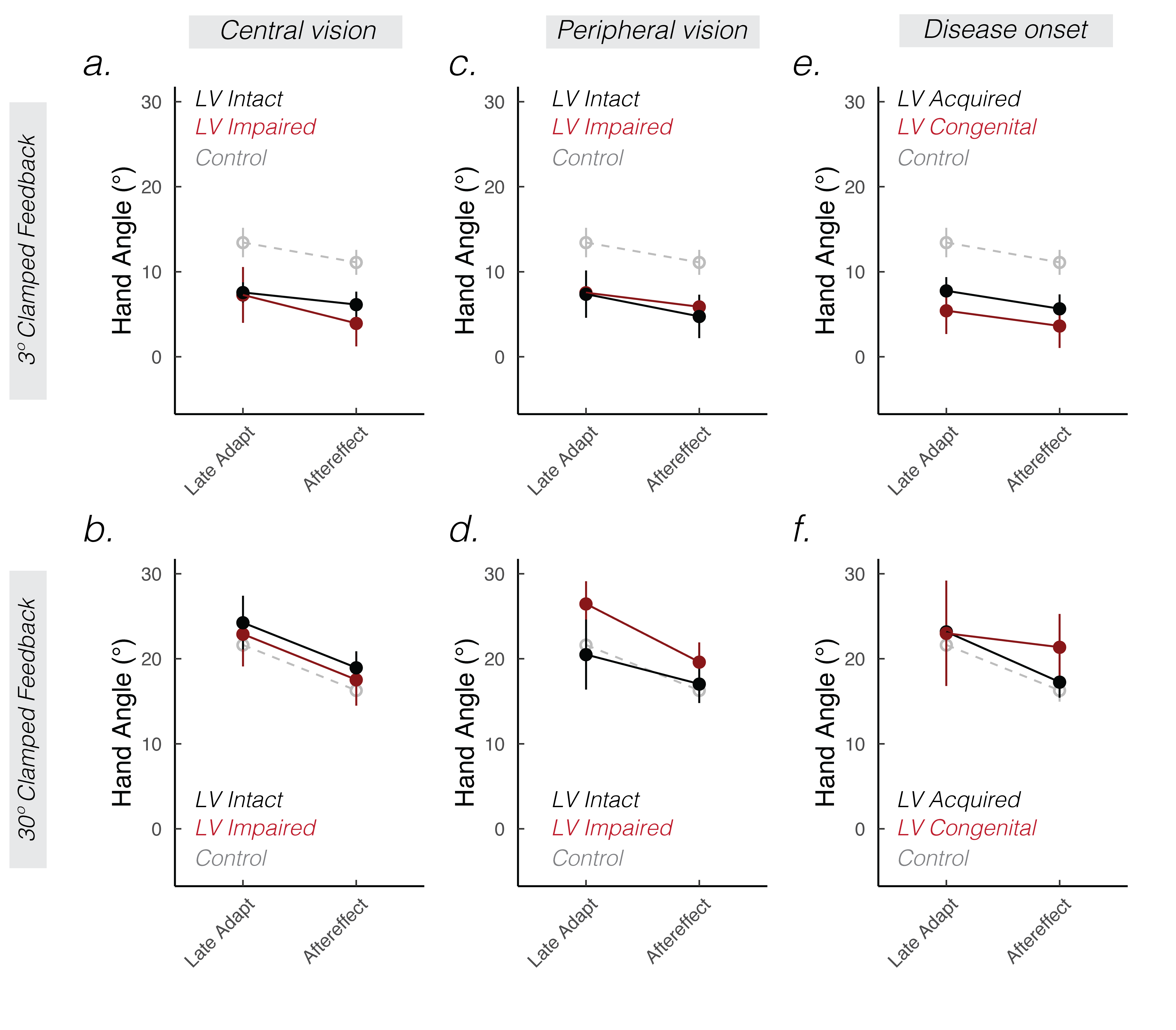

**Figure S1:** **Low vision subgroup analyses.** Mean hand angles ± SEM during late adaptation and aftereffect phases. Each column divides the low vision group based on a different performance or clinical variable: central vision loss **(a, b)**, peripheral vision loss **(c, d)**, or disease onset **(e, f)**. The control group is shown in grey dashed lines. * denotes p < 0.05. ^#^ denotes p < 0.10.

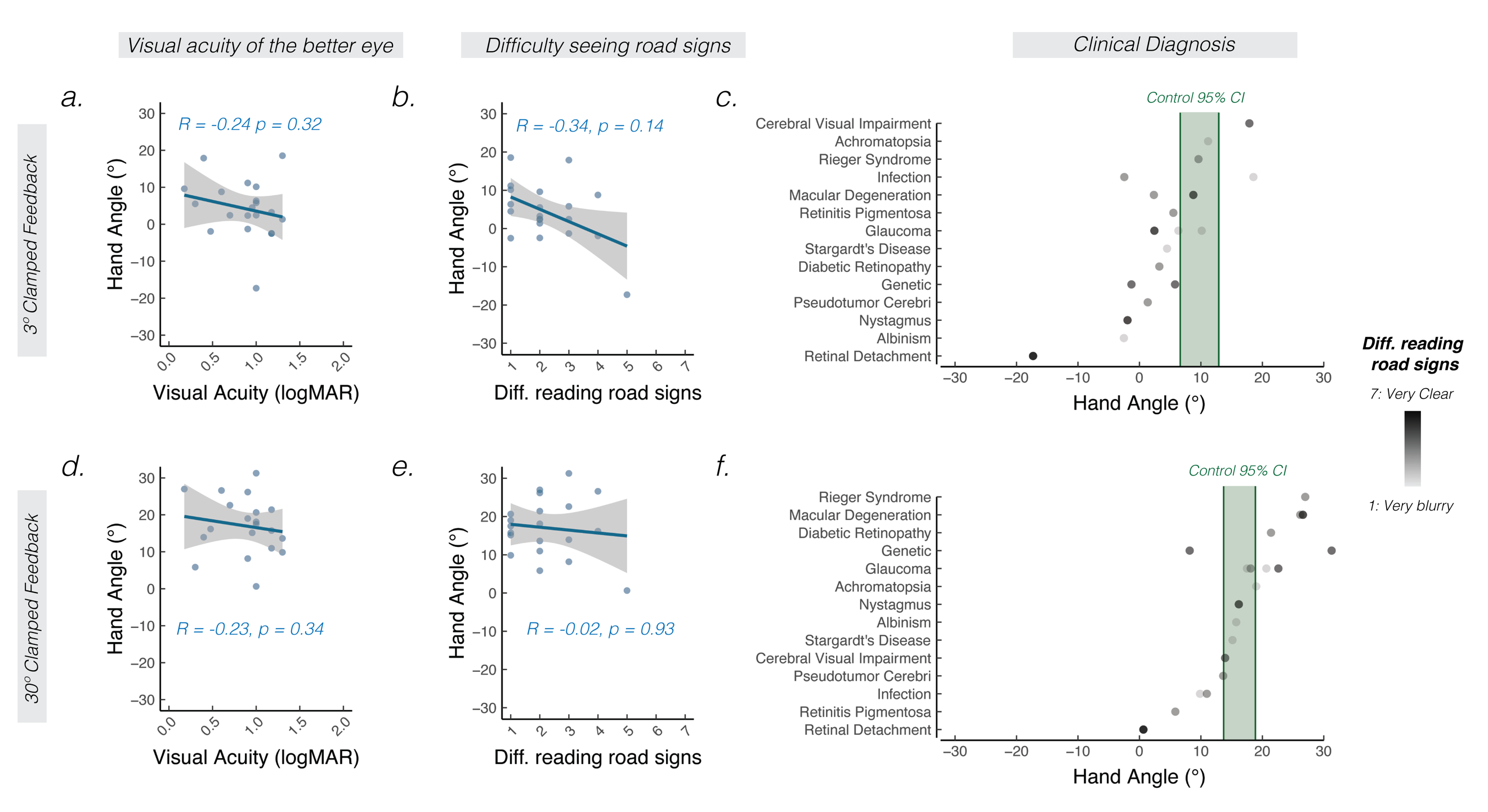

**Figure S2:** **The effect of visual acuity and clinical diagnosis on motor aftereffects.** Correlation between visual acuity of the less impaired eye and motor aftereffects **(a, d)**. Correlation between how clearly participants report seeing road signs (1 = very clear; 7 = very blurry) and motor aftereffects **(b, e).** Solid line indicates mean regression line and shaded region indicates SEM. The Spearman correlation is noted by *R*. **(c, f)** Mean aftereffects sorted by clinical diagnoses involving low vision. Shading of the dot indicates how well participants report seeing road signs (light shading = road signs are very blurry; dark shading = road signs are very clear). The 95% confidence interval for the control group is indicated by the green shaded region.

**Supplemental Tables**

| **#** | **Age** | **YOE** | **Hand** | **Gender** | **Low vision**  **diagnosis** | **Visual acuity in better eye** | **Seeing road signs** | **Peripheral visual field** | **Central**  **visual field** | **Low vision onset** |
| --- | --- | --- | --- | --- | --- | --- | --- | --- | --- | --- |
| 1 | 31 | 18 | R | M | Retinitis Pigmentosa | 0.3 | 2 | $\boldsymbol{\times}$ | **✓** | A |
| 2 | 34 | 22 | R | M | Achromatopsia | 0.9 | 2 | **✓** | **✓** | C |
| 3 | 60 | 18 | R | D | Retinitis Pigmentosa | 1.3 | 1 | $\boldsymbol{\times}$ | $\mathbf{✓}$ | A |
| 4 | 86 | 16 | R | M | Glaucoma | 1.0 | 2 | $\boldsymbol{\times}$ | **✓** | A |
| 5 | 70 | 18 | R | F | Macular Degeneration | 0.9 | 5 | **✓** | $\boldsymbol{\times}$ | A |
| 6 | 34 | 14 | R | F | Pseudotumor Cerebri | 1.7 | 1 | $\boldsymbol{\times}$ | $\boldsymbol{\times}$ | A |
| 7 | 61 | 18 | A | F | Rieger Syndrome | 0.2 | 4 | $\boldsymbol{\times}$ | **✓** | A |
| 8 | 26 | 16 | R | F | Charcot Marie Tooth | 0.9 | 4 | **✓** | **✓** | A |
| 9 | 86 | 16 | R | M | Glaucoma | 1 | 1 | $\boldsymbol{\times}$ | **✓** | A |
| 10 | 55 | 16 | R | M | Stargardt's Disease | 1 | 1 | $\boldsymbol{\times}$ | $\boldsymbol{\times}$ | A |
| 11 | 37 | 22 | A | F | Retinal Detachment | 1 | 5 | **✓** | **✓** | A |
| 12 | 29 | 18 | R | M | Glaucoma | 0.7 | 3 | $\boldsymbol{\times}$ | **✓** | A |
| 13 | 59 | 16 | R | M | Diabetic Retinopathy | 1.7 | 2 | $\boldsymbol{\times}$ | **✓** | A |
| 14 | 31 | 12 | R | F | Glaucoma, Deprivation Amblyopia | 1 | 1 | $\boldsymbol{\times}$ | **✓** | C |
| 15 | 33 | 18 | R | F | Genetic/Developmental | 1 | 3 | **✓** | $\boldsymbol{\times}$ | C |
| 16 | 24 | 16 | R | F | Albinism | 1.7 | 1 | $\boldsymbol{\times}$ | $\boldsymbol{\times}$ | C |
| 17 | 78 | 22 | R | F | Macular Degeneration | 0.6 | 4 | **✓** | $\boldsymbol{\times}$ | A |
| 18 | 28 | 18 | R | F | Nystagmus | 0.5 | 4 | **✓** | **✓** | C |
| 19 | 60 | 18 | A | F | Cerebral Visual Impairment | 0.4 | 3 | $\boldsymbol{\times}$ | **✓** | A |
| 20 | 60 | 18 | R | F | Infection | 1.7 | 2 | **✓** | $\boldsymbol{\times}$ | C |

**Table S1: Participants with low vision.** Gender was self-reported as male (M), female (F), or declined to specify (D). Handedness was reported as right (R), left (L), or unknown (U). Self-reports of visual acuity of the better seeing eye (logMAR) and peripheral/central visual field loss are provided. ✓ denotes intact and $\times$ denotes impaired function. Disease onset was self-reported as congenital (C), acquired (A), or unknown (U).
